# Leveraging AI for facioscapulohumeral muscular dystrophy prediction and omics biomarker identification

**DOI:** 10.1101/2025.10.01.679831

**Authors:** Joung Min Choi, Yi-Wen Chen, Liqing Zhang

## Abstract

Facioscapulohumeral muscular dystrophy (FSHD) is an autosomal dominant muscle disorder characterized by a complex genetic etiology, variable prognosis, and a lack of effective therapies. Previous studies have identified candidate protein and miRNA biomarkers using various profiling techniques, underscoring their potential for monitoring FSHD, assessing prognosis, and evaluating pharmacodynamic responses. However, the feasibility of applying machine learning (ML) models to predict FSHD using these molecular signatures has not been explored. In this study, we developed ML models to predict FSHD using a multi-omics dataset comprising protein abundance and miRNA expression profiles. Key predictive features were identified using Random Forest and the Support Vector Machine-Recursive Feature Elimination (SVM-RFE) methods. Performance evaluations demonstrated the robustness of the ML classifiers, with logistic regression consistently achieving the robust predictive accuracy in distinguishing FSHD from healthy conditions. Additionally, we assessed the predictive power of the identified features by comparing them with biomarker sets reported in previous studies. Our findings highlight the potential of AI to improve prediction accuracy and facilitate the cost- and time-efficient strategy for identifying FSHD biomarker candidates, even with limited sample sizes in the context of rare diseases.

## 1. Introduction

Facioscapulohumeral muscular dystrophy (FSHD) is an autosomal dominant muscle disorder characterized by variable prognosis, complex genetic and molecular mechanisms, and the absence of an effective therapy [1]. As one of the most common muscular dystrophies, it affects approximately 38,000 individuals in the United States and 870,000 worldwide [2, 3]. FSHD is primarily caused by the aberrant expression of double homeobox 4 (DUX4), resulting from epigenetic changes in the D4Z4 repeat region on chromosome 4q35 [4, 2, 3]. While no effective treatments are currently available, several promising therapeutic approaches are under active investigation.

FSHD is a late-onset disease, with approximately 4% of cases presenting early onset [5]. Most patients are diagnosed during their late teens or adult years [6, 7, 8]. The causative gene, DUX4, is expressed stochastically and at extremely low levels, making its detection in patient muscle tissue particularly challenging [9, 10, 11]. Recent studies suggest that magnetic resonance imaging (MRI)-guided biopsies from muscles exhibiting active disease activity improve the likelihood of detecting DUX4 mRNA [12, 13]. However, even with this approach, DUX4 expression remains low and may be undetectable in certain samples. Functional motor scales offer a non-invasive alternative to monitor neuromuscular disease progression but are limited by variability, dependence on age or disease stage, and susceptibility to placebo or coaching effects in clinical trials [14, 15]. MRI has proven valuable for identifying affected muscles and regions with active disease activity and pathological changes [16]. Nevertheless, its high cost, requirement for specialized equipment, time-intensive nature, and inability to detect subtle or early molecular changes pose significant challenges. The slow progression of FSHD further complicates the evaluation of treatment efficacy using motor function data or imaging findings.

To enhance clinical management and facilitate the development of new treatments, non-invasive methods for detecting and studying FSHD are essential. Circulating molecular biomarkers offer a promising alternative to traditional clinical assessments, providing objective measurements that can be repeatedly assayed over time using minimally invasive techniques [17, 18].

Additionally, these biomarkers may respond rapidly to treatments that improve the disease, offering real-time insights into therapeutic efficacy. Blood-based miRNAs or proteins that change with disease progression or in response to therapy are categorized as monitoring and pharmacodynamic biomarkers, respectively [19]. In clinical trials, monitoring biomarkers can serve as treatment-response biomarkers, helping identify therapy responders, establish treatment-response relationships, and improve statistical power and modeling. Therefore, the development of less invasive monitoring and pharmacodynamic biomarkers is critical for advancing FSHD therapeutics. Previously, Statland et al. identified seven protein biomarker candidates by analyzing 22 FSHD serum samples using a commercial multiplex assay [20]. A multi-site study employing SomaScan proteomics in two independent FSHD populations identified four proteins that showed consistent behavior across cohorts [21]. To broaden the scope beyond proteins pre-selected by specific platforms, proteomics profiling was performed to identify proteins that differ in FSHD plasma samples compared to age- and sex-matched healthy controls. Additionally, circulating miRNAs were examined using TaqMan miRNA arrays to identify differentially expressed miRNAs [22]. In this study, a total of 32 miRNAs and proteins were statistically identified as significantly altered in the plasma of individuals with FSHD compared to healthy controls. These findings underscore the potential of circulating biomarkers to revolutionize FSHD research and therapeutic development.

Recently, the integration of molecular profile datasets with machine learning (ML), a prominent artificial intelligence (AI) technique, has gained widespread adoption in disease prediction, particularly in oncology, where it has demonstrated high accuracy in prediction and identification of key molecular features [23, 24, 25, 26, 27]. Compared to traditional histological assessments that require biopsies, applying ML models to molecular profiles offers faster, more efficient, and more accurate prediction tools.

In this study, we applied machine learning models to predict FSHD using a multi-omics dataset comprising protein abundance and miRNA expression profiles. Widely used classification models, including Support Vector Machine (SVM), Logistic Regression (LR), and Random Forest (RF), were employed to classify FSHD and identify key predictive features. These features, which hold potential as biomarkers, were identified using feature importance scores from RF and the Support Vector Machine-Recursive Feature Elimination (SVM-RFE) method. Performance evaluation experiments confirmed the robustness and stability of these classifiers, demonstrating the strong predictive power of the selected feature sets. This approach offers a promising enhancement in differentiating FSHD from healthy conditions. Further investigations into the identified feature sets highlight the efficacy and potential of AI as a reliable method for identifying biomarker candidates for FSHD prediction, even in the context of rare diseases and limited sample sizes.

## 2. Methods

### 2.1. Dataset Collection

The dataset was obtained from Heier et al. [22] and includes two types of omics data: proteomics plasma samples and miRNA profiles. The data were collected from FSHD patients aged 10 to 51 years and healthy control volunteers aged 16 to 54 years. According to the dataset description, all FSHD patients had Type 1 FSHD, characterized by epigenetic changes due to D4Z4 contraction, which leads to the upregulation of DUX4. Among the participants, 13 FSHD patients and 8 healthy controls provided multi-omics profiles, comprising both proteomics and miRNA data, as summarized in Table 1.

**Table 1:** Dataset used for evaluating the ML classifiers for FSHD prediction.

| Omics type | Proteomics | miRNA | Multi-omics |
| --- | --- | --- | --- |
| # of FSHD samples | 25 | 16 | 13 |
| # of control samples | 16 | 8 | 8 |

### 2.2. Preprocessing

For the proteomics plasma samples, features with identical unique peptide counts across all samples were removed, and only the features common to all samples were retained. Then, min-max normalization was applied. For the miRNA dataset, “undetermined” values were first set to 40 following the preprocessing step described in the original study [22]. The dataset then underwent a preprocessing approach similar to that used for the proteomics profiles. After preprocessing, 150 protein features and 258 miRNA features remained.

### 2.3. FSHD prediction using ML classification models

For FSHD prediction, we employed three widely-used ML classification models: Support Vector Machine (SVM) with a linear kernel, Logistic Regression (LR), and Random Forest (RF). SVM is a supervised learning algorithm that identifies a hyperplane to create a decision boundary, classifying data points by maximizing the margin between classes. RF, an extension of the decision tree, constructs a tree-structured classification model based on a set of discrete rules from the training data. Starting at the root node, RF follows the appropriate branches according to the decision rules, assigning data points to classes based on the terminal node. RF is an ensemble learning method that aggregates the outputs from multiple decision trees (DTs) to derive a final prediction. LR estimates the probability of class membership by applying a sigmoid function to the output of a linear regression function. These three models were chosen due to their strong classification performance compared to alternatives like Naive Bayes and individual decision trees [28]. The classifiers were implemented using the “Scikit-learn” Python package [29] with default parameter settings.

### 2.4. Identification of biomarker candidates for FSHD prediction

To identify the features that contributed most to FSHD prediction, and that might serve as potential biomarkers, we measured feature importance using RF and Support Vector Machine-Recursive Feature Elimination (SVM-RFE). SVM-RFE iteratively trains a SVM model, ranks features based on their contribution to the decision boundary, and assigns ranks where lower-ranked features are less important. In each iteration, the least significant features are removed, and this process continues until a specified number of features remain. From the results of both methods, we selected the top 50 most important features from each and then extracted the features shared by both approaches. This analysis was applied to both proteomics and miRNA profiles, resulting in the identification of 20 protein features and 15 miRNAs.

## 3. Results and Discussion

### 3.1. FSHD prediction improvement by multi-omics dataset

To evaluate whether utilizing multi-omics datasets could improve the prediction of FSHD, we compared the performance of ML classification models trained on multi-omics versus single-omics data. A 5-fold cross-validation was conducted. In this experiment, LR and SVM demonstrated improved performance when trained on multi-omics datasets compared to single-omics datasets (Table 2, Fig 1). The multi-omics dataset, which combined proteomics and miRNA expression profiles, achieved the highest average accuracy of 0.700 and an F1-score of 0.785 using LR. In contrast, RF showed a slight decrease in prediction performance when using the multi-omics dataset, with the average F1-score dropping from 0.735 (proteomics) to 0.651 (multiomics). This decline may be attributed to the reduced number of samples in the multi-omics dataset (21 samples) compared to the proteomics dataset (41 samples).

**Table 2:** Average classification performance with 95% confidence interval of ML classifiers for FSHD prediction using different omics datasets based on 5-fold cross validation on Heier et al. dataset.

| Model | Metric | Multi-omics | Single-omics (Proteomics) | Single-omics (miRNA) |
| --- | --- | --- | --- | --- |
| LR | Accuracy | 0.700±0.139 | 0.611±0.116 | 0.670±0.213 |
| | F1-score | 0.785±0.088 | 0.724±0.122 | 0.779±0.155 |
| | Precision | 0.750±0.179 | 0.639±0.137 | 0.750±0.323 |
| | Recall | 0.867±0.227 | 0.900±0.278 | 0.900±0.170 |
| SVM | Accuracy | 0.670±0.213 | 0.564±0.107 | 0.650±0.170 |
| | F1-score | 0.779±0.155 | 0.682±0.128 | 0.725±0.179 |
| | Precision | 0.750±0.323 | 0.618±0.158 | 0.717±0.227 |
| | Recall | 0.900±0.170 | 0.817±0.258 | 0.767±0.278 |
| RF | Accuracy | 0.550±0.260 | 0.686±0.270 | 0.540±0.111 |
| | F1-score | 0.645±0.241 | 0.735±0.258 | 0.681±0.092 |
| | Precision | 0.683±0.296 | 0.705±0.181 | 0.720±0.333 |
| | Recall | 0.700±0.370 | 0.817±0.361 | 0.800±0.340 |

**Figure 1.**
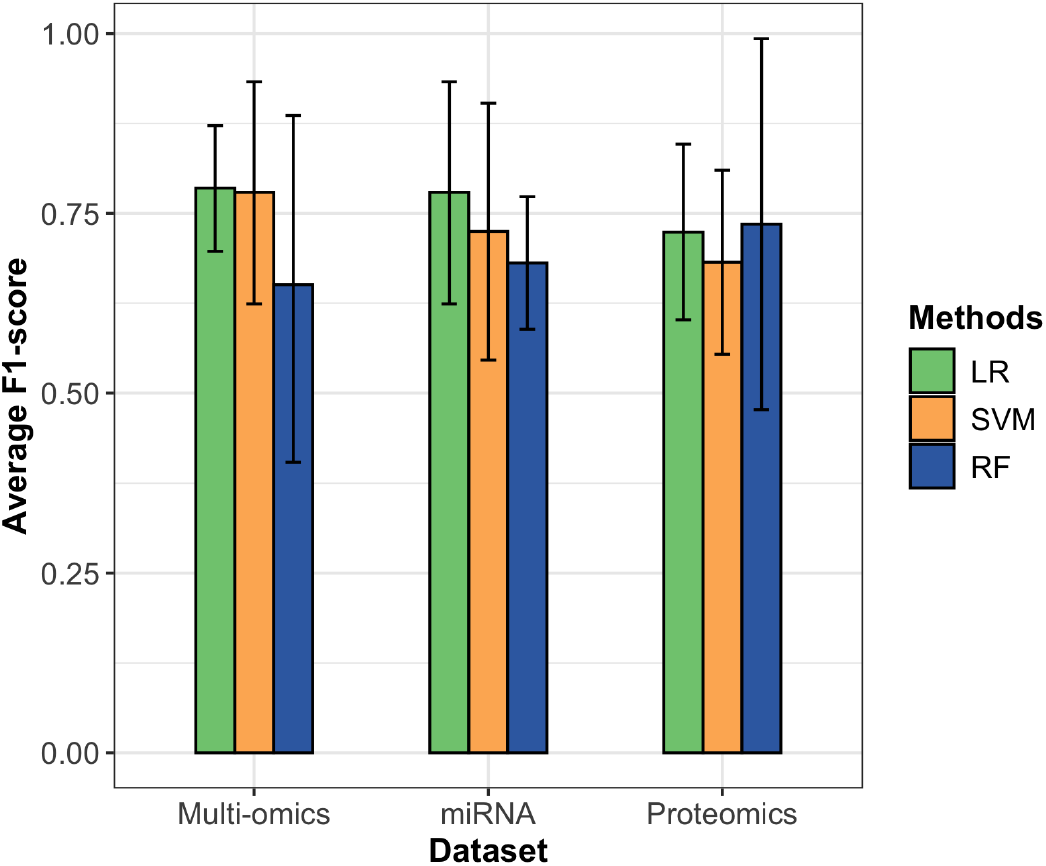
Performance comparison of ML classifiers in FSHD prediction using different omics dataset based on average F1-score with 95% confidence interval from 5-fold cross validation results.

### 3.2. Investigation of biomarker candidates for FSHD prediction

The features that contributed most to FSHD prediction have the potential to serve as biomarkers for the disease. We evaluated the importance scores of each feature across different omics profiles and selected the top 50 features using RF and SVM-RFE. We then identified the overlapping features between the two methods. As a result, 20 protein features and 15 miRNAs were identified (Supplementary Material S1). To assess the potential of these features as biomarker candidates, we conducted several analyses. First, to directly compare the abundance differences between FSHD patients and control groups, we visualized the normalized expression of the miRNAs and protein abundance using the Heier et al. dataset. A t-test was performed to determine the statistical differences between the FSHD and control groups and fold changes were calculated based on the average normalized expression of miRNAs and the average unique peptide counts of protein features. Based on the results (Fig. 2, Supplementary Material S2), seven out of the 15 miRNA features (hsa-miR-103, hsa-miR-29, hsa-miR-98, hsa-miR-505, has-miR-34, hsa-miR-329, hsa-miR-9) showed significant differences between the two groups (p-value *<* 0.05). Several of these miRNAs have been previously implicated in muscle pathology relevant to FSHD. Notably, hsa-miR-29b has been shown to modulate TGF-*β* signaling, a key pathway involved in muscle fibrosis and atrophy—hallmarks of FSHD progression—particularly through the regulation of myostatin, a negative regulator of muscle growth [30, 31, 18]. hsa-miR-98 has also been demonstrated to inhibit TGF-*β* and pro-inflammatory cytokines, suggesting its potential involvement in FSHD by modulating muscle inflammation and fibrotic processes [32]. hsa-miR-505 has been associated with muscle regeneration and cardiac muscle remodeling, suggesting a broader role in muscle tissue homeostasis [33]; its dysregulation in FSHD may reflect underlying muscle remodeling processes such as impaired regeneration and fibrosis driven by aberrant DUX4 expression. hsa-miR-9, which is known to inhibit satellite cell activation, may contribute to impaired muscle regeneration—a known feature of FSHD muscle pathology [34, 35]. The observed dysregulation of these miRNAs in FSHD patient samples supports their potential relevance to the disease and highlights them as promising biomarker candidates.

**Figure 2.**
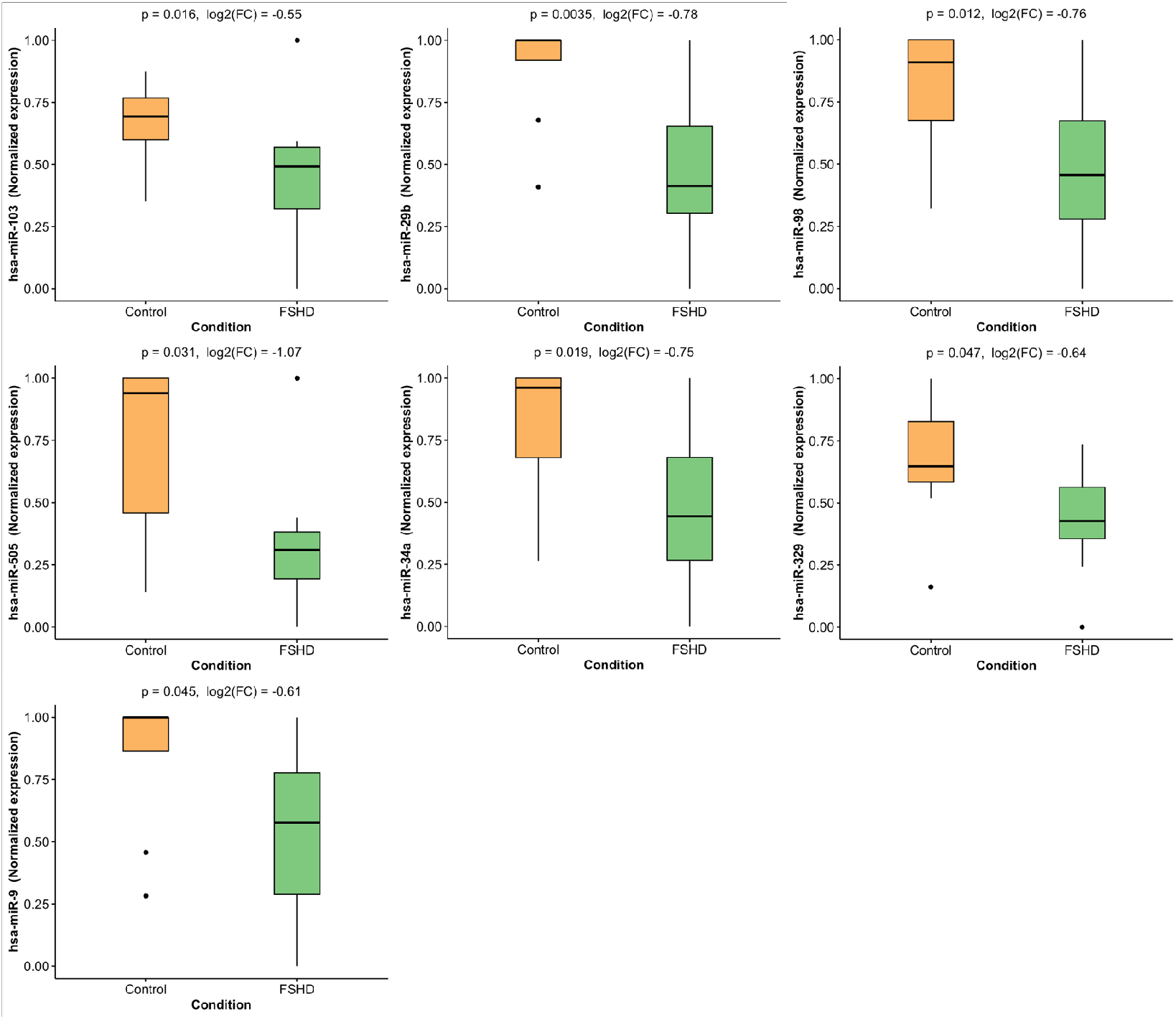
Normalized expression values of the top miRNAs with the highest importance scores for FSHD prediction, based on the Heier et al. dataset, showing a statistically significant difference (p-value < 0.05) between control and FSHD patients.

For the protein features, while the results were not statistically significant, differences in abundance between the two groups were observed (Fig. 3, Supplementary Material S3). Among the 20 features analyzed, HP and PF4 exhibited the highest fold changes in normalized peptide counts in FSHD patients compared to healthy controls. HP has been linked to inflammatory and oxidative stress responses in neuromuscular diseases such as Duchenne muscular dystrophy [36], while PF4 modulates immune signaling [37], both of which may relate to immune dysregulation in FSHD muscle. We also evaluated biomarker sets from four prior studies that identified protein biomarkers for FSHD: Statland et al. [20], Petek et al. [21], Corosolla et al. [38], and Wong et al. [39]. These datasets were compared with the proteins identified as important in our analysis to identify common features. VTN (Vitronectin), reported to be downregulated in FSHD muscles in previous studies [20, 38], consistently with an overall downregulation of the plasminogen pathway, which also aligns with our findings (Fig. 3). Similarly, AGT (Angiotensinogen), GSN (Gelsolin), and HPX (Hemopexin), which have been reported as having altered regulation in FSHD [20, 38], showed consistent patterns in our analysis. AGT promotes fibrosis and inflammation via the angiotensin/TGF-*β*1 axis [40]; GSN regulates actin filament turnover, crucial for muscle repair [41]; and HPX contributes to heme scavenging, potentially mitigating oxidative stress—all of which are relevant to key pathological pathways in muscle. C5 (Complement C5) was noted by Wong et al. to exhibit a trend toward elevation in FSHD samples compared to controls [39], a finding corroborated by our results. Wong et al. further reported that human skeletal muscle cells with DUX4 expression induce RNAs for several complement components, including C5. They also highlighted that monoclonal antibodies targeting C5 are used to treat conditions such as myasthenia gravis, paroxysmal nocturnal hemoglobinuria, atypical hemolytic uremic syndrome, and neuromyelitis optica [39]. Lastly, CFI (Complement Factor I), identified as a potential biomarker due to its elevation in FSHD [21, 39], also demonstrated a similar trend in our analysis.

**Figure 3.**
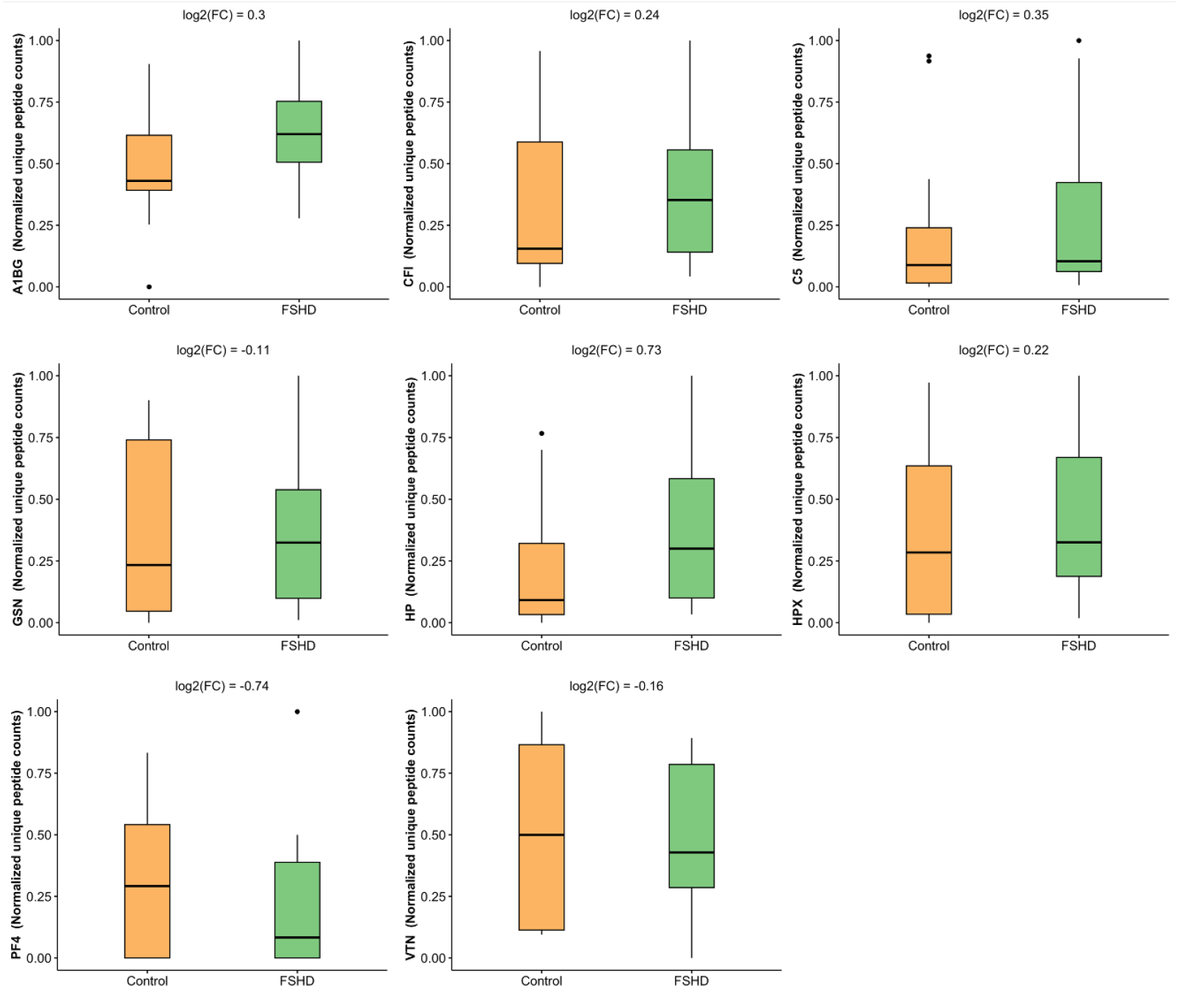
Normalized unique peptide counts of the top proteins with the highest importance for FSHD prediction, based on the Heier et al. dataset.

### 3.3. Exploring the predictive power of biomarker candidates for FSHD prediction

We tested the predictive power of these identified features for FSHD prediction. ML classifiers were trained using only the important features and compared to classifiers trained with all features to determine if similar or improved performance could be achieved. First, 5-fold cross-validation was conducted using the Heier et al. dataset across both single-omics and multiomics profiles. The results (Fig. 4, Table 3) showed that when ML models were trained using only the important features, the FSHD prediction performance improved across all omics datasets, except for the LR model for miRNA, which yielded a comparable average F1-score (0.740 with important features vs. 0.779 with all features). Notably, for the multi-omics dataset, using the important feature set significantly improved the average F1-score: from 0.779 to 0.943 for SVM, from 0.651 to 0.731 for RF, and from 0.785 to 0.836 for LR. Across the proteomics dataset, all three ML classifiers demonstrated improvement, with RF achieving the highest performance, marked by an average accuracy of 0.761 and an F1-score of 0.780, compared to its previous scores of 0.686 and 0.735, respectively.

**Table 3:** Average classification performance with 95% confidence interval of ML classifiers for FSHD prediction using different feature sets for model training across various omics datasets based on 5-fold cross-validation. (‘imp’ denotes using identified important feature set and ‘all’ denotes using all features)

|  | Method | SVM |  | RF |  | LR |  |
| --- | --- | --- | --- | --- | --- | --- | --- |
| Omics | Feature set | All set | Imp set | All set | Imp set | All set | Imp set |
| Multi-omics | Accuracy | 0.670±0.213 | **0.900**±0.170 | 0.550±0.260 | **0.650**±0.170 | 0.700±0.139 | **0.750**±0.219 |
| | F1-score | 0.779±0.155 | **0.943**±0.097 | 0.645±0.241 | **0.731**±0.113 | 0.785±0.088 | **0.836**±0.149 |
| Proteomics | Accuracy | 0.564±0.107 | **0.636**±0.087 | 0.686±0.270 | **0.761**±0.273 | 0.611±0.116 | **0.664**±0.199 |
| | F1-score | 0.682±0.128 | **0.737**±0.099 | 0.735±0.258 | **0.780**±0.269 | 0.724±0.122 | **0.776**±0.142 |
| miRNA | Accuracy | 0.650±0.170 | **0.710**±0.127 | 0.540±0.111 | **0.800**±0.304 | **0.670**±0.213 | 0.630±0.194 |
| | F1-score | 0.725±0.179 | **0.786**±0.100 | 0.681±0.092 | **0.846**±0.220 | **0.779**±0.155 | 0.740±0.154 |

**Figure 4.**
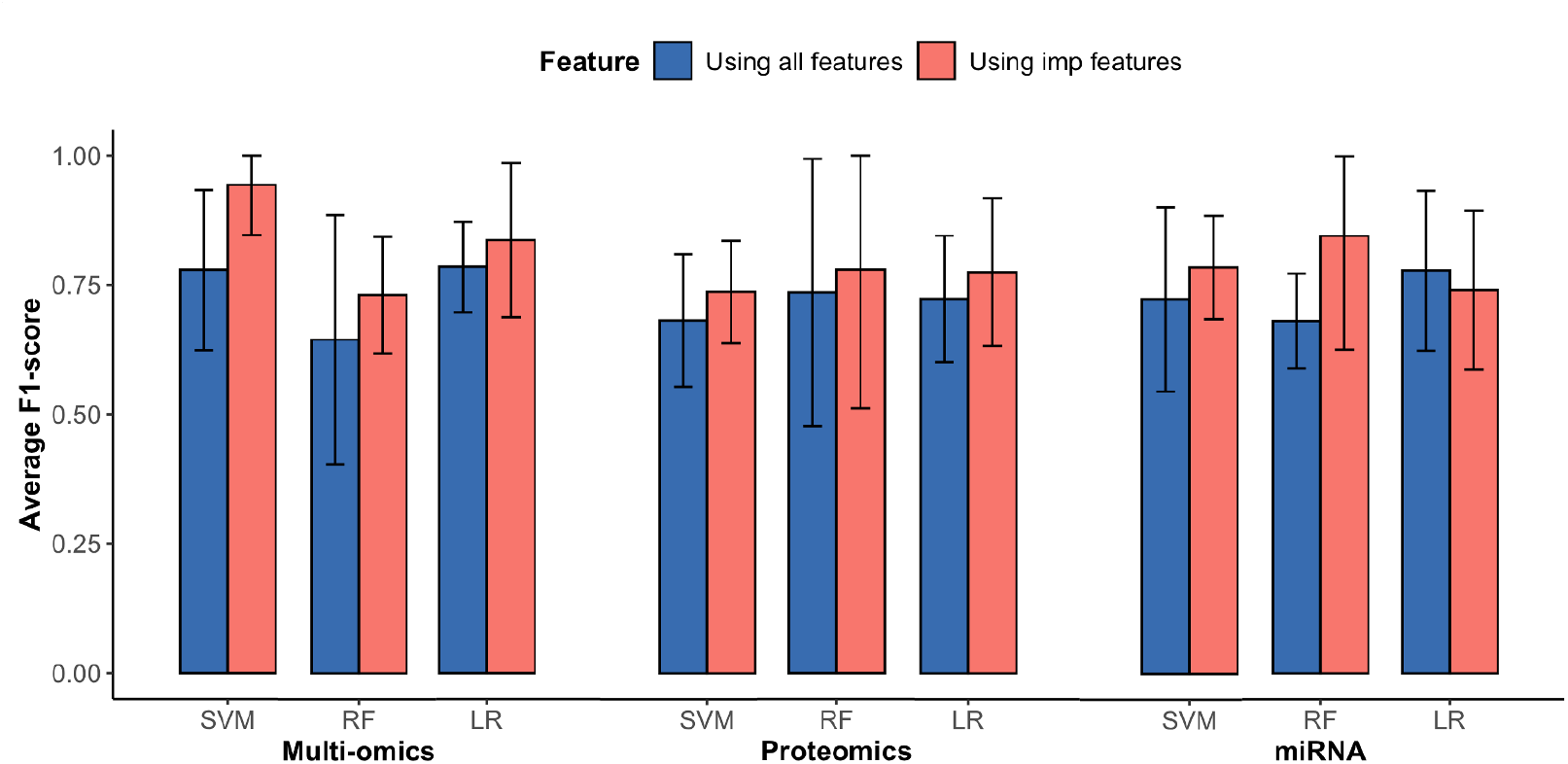
Performance improvement in FSHD prediction using ML classification models trained with identified important features across omics datasets based on 5-fold crossvalidation. 95% confidence interval is denoted with error bars.

### 3.4. Evaluating prediction performance using biomarker candidates identified by ML and prior studies

To further validate the predictive power of the identified biomarker candidates, we compared the performance of ML classifiers trained using these biomarkers against those trained with biomarker sets reported in previous studies. Five studies that identified protein biomarkers for FSHD were selected [20, 21, 22, 38, 39], and their biomarker lists were cross-referenced with our dataset. The number of overlapping biomarkers used for model training was as follows: Statland et al. (5), Petek et al. (6), Corasolla et al. (27), and Wong et al. (10). ML model performance was evaluated using five-fold cross-validation. The results, summarized in Fig. 5, Table 4, and Supplementary Material S4, indicate that the important features identified by ML yielded comparable or superior prediction performance to the biomarker sets provided by the other studies. For LR, the ML-identified features achieved the highest average F1-score of 0.776, with the Corasolla et al. biomarker set closely following at 0.766. Similarly, for RF and SVM, the ML-identified features attained F1-scores of 0.780 and 0.737, respectively, while the Corasolla et al. biomarker set showed equivalent performance for RF (0.780) and a slightly higher score for SVM (0.759). These findings demonstrate that the feature sets identified by ML possess robust predictive power, comparable to or better than existing biomarker sets. This underscores their potential utility as reliable biomarker candidates to distinguish FSHD from healthy conditions.

**Table 4:**
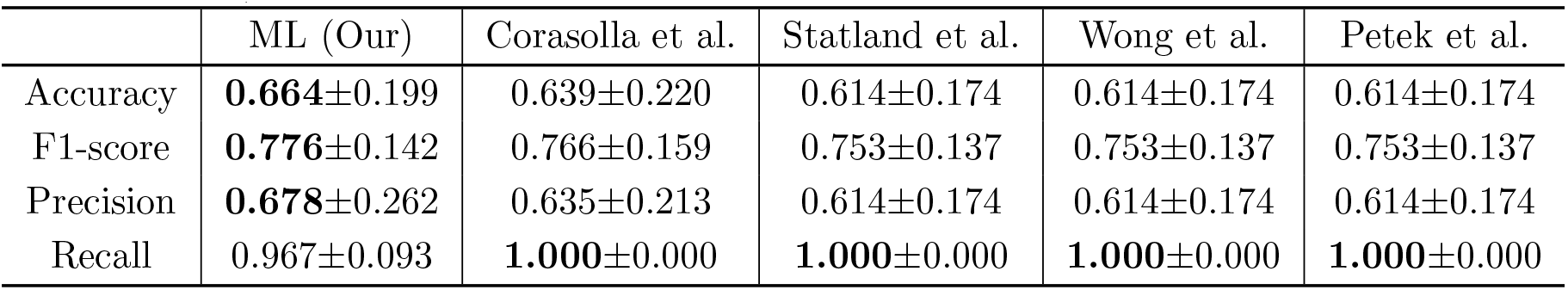
Average classification performance of the LR model for FSHD prediction using biomarker sets, evaluated via five-fold cross-validation.

|  | ML (Our) | Corasolla et al. | Statland et al. | Wong et al. | Petek et al. |
| --- | --- | --- | --- | --- | --- |
| Accuracy | **0.664**±0.199 | 0.639±0.220 | 0.614±0.174 | 0.614±0.174 | 0.614±0.174 |
| F1-score | **0.776**±0.142 | 0.766±0.159 | 0.753±0.137 | 0.753±0.137 | 0.753±0.137 |
| Precision | **0.678**±0.262 | 0.635±0.213 | 0.614±0.174 | 0.614±0.174 | 0.614±0.174 |
| Recall | 0.967±0.093 | **1.000**±0.000 | **1.000**±0.000 | **1.000**±0.000 | **1.000**±0.000 |

**Figure 5.**
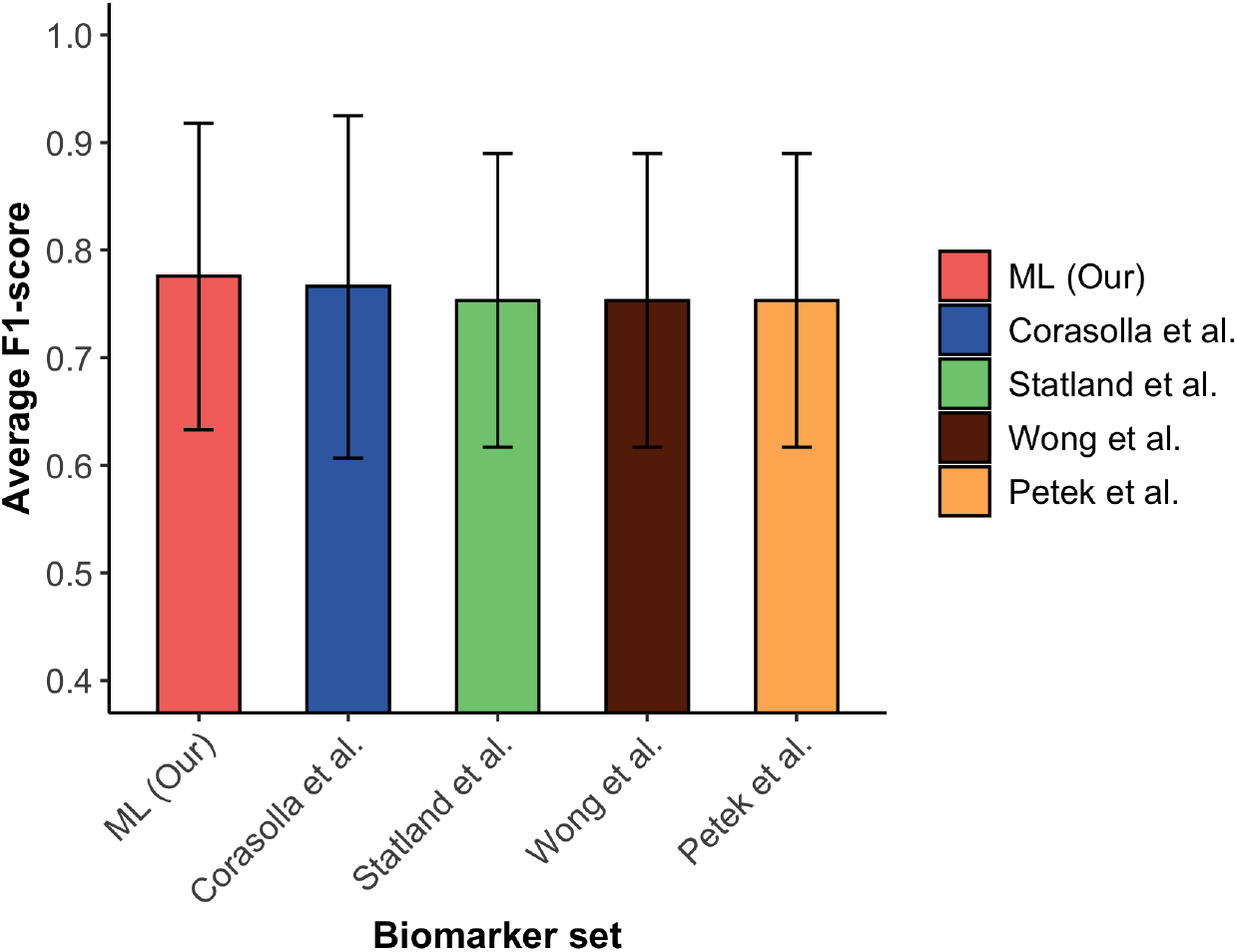
Comparison of FSHD prediction performance based on average F1-scores of the LR model trained with various biomarker sets, using five-fold cross-validation.

### 3.5. Assessing FSHD prediction performance with varying numbers of biomarkers

To provide researchers with guidance on selecting the optimal number of biomarkers for FSHD prediction, we investigated how prediction performance changes as the number of biomarkers varies. In each iteration, the least important biomarker was sequentially removed, and five-fold cross-validation was conducted to measure the average performance using SVM, RF, and LR. This process continued until only one feature remained. The ranked lists of 20 protein features and 15 miRNA biomarker candidates are provided in Supplementary Material S1. The results, summarized in Fig. 6 and Supplementary Material S5, indicate that in the proteomics dataset, training SVM and RF models with 14 protein markers resulted in the highest average F1-scores (0.754 and 0.798, respectively), compared to using all 20 protein features, which yielded F1-scores of 0.737 and 0.789. LR performance remained consistent, achieving an F1-score of 0.776 with both 14 and 20 features. For the miRNA dataset, the SVM and LR models performed best with 7 miRNA markers, both achieving an average F1-score of 0.836, whereas RF showed its highest performance when using all 15 miRNA features. For the multi-omics dataset, reducing the feature set to 24 markers (12 proteins and 12 miRNAs) produced the best results across all three machine learning models. The average F1-scores were 0.943 for SVM, 0.865 for RF, and 0.903 for LR, demonstrating improved performance compared to using the full 35-feature set.

**Figure 6.**
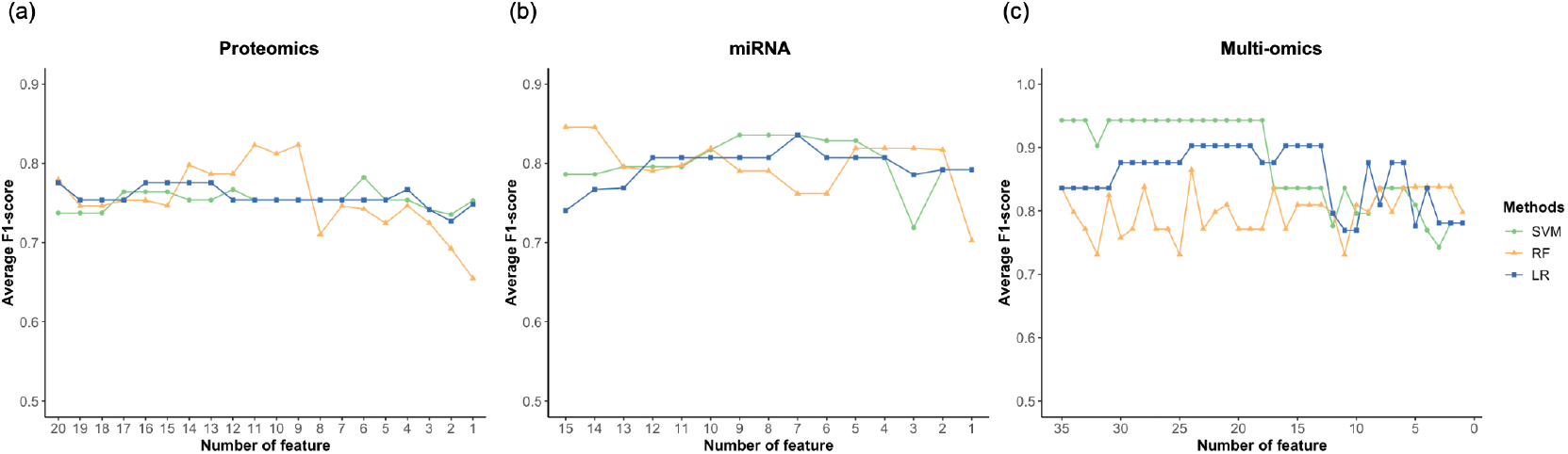
FSHD prediction performance across different numbers of biomarkers using (a) proteomics, (b) miRNA, and (c) multi-omics datasets. For each iteration, five-fold cross-validation was performed, progressively eliminating the least important feature for SVM, RF, and LR models, and the average F1-score was evaluated.

## 4. Conclusions

In this study, we demonstrated the application of ML-based models for FSHD prediction. Widely used ML classifiers, including SVM, RF, and LR, were implemented and evaluated using multi-omics datasets consisting of proteomics and miRNA profiles. Models trained on the multi-omics dataset generally achieved better prediction performance compared to those trained on single-omics datasets. Overall, LR showed better performance compared to other classifiers for FSHD prediction across different omics datasets.

We also identified key features contributing to FSHD prediction and assessed their potential as biomarkers for the disease. Notably, the selected fea-ture sets significantly enhanced predictive performance compared to models utilizing all features. Among the identified biomarkers, seven miRNAs exhibited statistically significant differences between FSHD and control groups and were consistent with findings from previous studies. Although protein features did not reach statistical significance, six proteins were identified as potential biomarkers, aligning with findings from other studies and showing similar patterns of alteration in FSHD cases compared to controls.

To validate the predictive power of the identified biomarker candidates, we compared the performance of ML classifiers trained using these biomarkers against those trained with biomarker sets reported in prior studies. The LR model trained on the identified biomarkers achieved the highest predictive performance, often surpassing or matching the performance of existing biomarker sets in other classifiers. These results highlight the potential of AI-driven approaches in identifying robust biomarkers to distinguish FSHD from healthy controls.

Additionally, we explored the impact of biomarker set size on prediction performance, offering guidance for researchers on optimal marker selection. For single-omics datasets, using 14 protein markers or 7 miRNA markers provided comparable or superior performance relative to larger sets. For multi-omics datasets, reducing the feature set to 24 markers (12 proteins and 12 miRNAs) yielded the best results across all three ML models. These experimental results could guide a way for researchers to predict FSHD more efficiently by reducing costs, for instance, through the use of targeted sequencing techniques.

FSHD is a rare genetic muscle disease, which can be underdiagnosed due to heterogenous presentation and technically-challenging genetic testing. The rarity of the disease presents challenges in acquiring large cohorts. Nonetheless, our findings demonstrate that ML classifiers can effectively identify biomarker candidates and yield robust prediction performance even with limited sample sizes. This underscores the value of AI in advancing FSHD research by enhancing prediction accuracy and enabling the efficient identification of biomarker candidates for differentiating FSHD from healthy conditions in a cost- and time-effective manner.

## Acknowledgements

Y.-W. C. is partially supported by FSHD Canada Foundation/SOLVE FSHD, NIH/NICHD 1R21HD103993-01, and NIH/NIAMS 1R21AR080887-01.LZ is partially supported by National Science Foundation (NSF) #2004751, #2125798, #2344169 and #2319522, as well as the National Institutes of Health (NIH) grant #1R01AI179686-01A1.

## Authors’ contributions

Joung Min Choi: Conceptualization, Methodology, Software, Investigation, Writing-Original Draft, Review & Editing, Visualization; Yi-Wen Chen: Writing-Review & Editing, Funding acquisition; Liqing Zhang: Writing-Review & Editing, Supervision, Project administration, Funding acquisition

## Ethical Publication Statement

We confirm that we have read the Journal’s position on issues involved in ethical publication and affirm that this report is consistent with those guidelines.

## Availability of data and materials

The data that support the findings of this study are openly available in https://www.mdpi.com/2075-4426/10/4/236.

