## Supplementary Material for "Leveraging AI for facioscapulohumeral muscular dystrophy prediction and omics biomarker identification"

**Supplementary Material S1**. The list of 20 protein features and 15 miRNAs identified as biomarker candidates using Random Forest (RF) and Support Vector Machine Recursive Feature Elimination (SVM-RFE). Feature importance scores across various omics profiles were measured, and the top 50 features were selected by both methods. The overlapping features between RF and SVM-RFE were identified as the final biomarker candidates.

Proteins

| Protein ID | Protein name | Gene name | RF ranking | SVM-RFE ranking |
|---|---|---|---|---|
| P04217 | Alpha-1B-glycoprotein | A1BG | 36 | 7 |
| P01019 | Angiotensinogen;Angiotensin-1;Angiotensin-2;Angiotensin-3;Angiotensin-4;Angiotensin 1-9;Angiotensin 1-7;Angiotensin 1-5;Angiotensin 1-4 | AGT | 12 | 43 |
| P02765 | Alpha-2-HS-glycoprotein;Alpha-2-HS-glycoprotein chain A;Alpha-2-HS-glycoprotein chain B | AHSG | 8 | 9 |
| P02647 | Apolipoprotein A-I;Proapolipoprotein A-I;Truncated apolipoprotein A-I | APOA1 | 27 | 17 |
| P02655 | Apolipoprotein C-II;Proapolipoprotein C-II | APOC2 | 26 | 1 |
| P01031 | Complement C5;Complement C5 beta chain;Complement C5 alpha chain;C5a anaphylatoxin;Complement C5 alpha chain | C5 | 20 | 32 |
| P16070 | CD44 antigen | CD44 | 29 | 3 |
| P05156 | Complement factor I;Complement factor I heavy chain;Complement factor I light chain | CFI | 35 | 39 |
| P03951 | Coagulation factor XI;Coagulation factor XIa heavy chain;\Coagulation factor XIa light chain | F11 | 16 | 28 |
| P06396 | Gelsolin | GSN | 18 | 6 |
| P00738 | Haptoglobin;Haptoglobin alpha chain;Haptoglobin beta chain | HP | 4 | 2 |
| P02790 | Hemopexin | HPX | 40 | 20 |
| P19823 | Inter-alpha-trypsin inhibitor heavy chain H2 | ITIH2 | 31 | 33 |
| P02750 | Leucine-rich alpha-2-glycoprotein | LRG1 | 17 | 10 |
| P51884 | Lumican | LUM | 38 | 47 |
| P02776 | Platelet factor 4;Platelet factor 4, short form | PF4 | 6 | 11 |
| P02775 | Platelet basic protein;Connective tissue-activating peptide III;TC-2;Connective tissue-activating peptide III(1-81);Beta-thromboglobulin; Neutrophil-activating peptide 2(74);Neutrophil-activating peptide 2(73);Neutrophil-activating peptide 2;TC-1;Neutrophil-activating peptide 2(1-66);Neutrophil-activating peptide 2(1-63) | PPBP | 5 | 29 |
| P01008 | Antithrombin-III | SERPINC1 | 21 | 35 |
| P02766 | Transthyretin | TTR | 2 | 27 |
| P04004 | Vitronectin;Vitronectin V65 subunit;Vitronectin V10 subunit;Somatomedin-B | VTN | 11 | 8 |

miRNAs

| miRNA | RF ranking | SVM-RFE ranking |
|---|---|---|
| hsa-miR-103 | 1 | 15 |
| hsa-miR-329 | 2 | 18 |
| hsa-miR-29b | 4 | 14 |
| hsa-miR-138 | 5 | 5 |
| hsa-miR-98 | 6 | 28 |
| hsa-miR-885-5p | 12 | 32 |
| hsa-miR-522 | 14 | 9 |
| hsa-miR-505 | 15 | 3 |
| hsa-miR-122 | 16 | 27 |
| hsa-miR-200c | 19 | 24 |
| hsa-miR-375 | 21 | 12 |
| hsa-miR-576-3p | 23 | 41 |
| hsa-miR-34a | 29 | 20 |
| hsa-miR-9 | 30 | 30 |
| hsa-miR-365 | 45 | 43 |

**Supplementary Material S2**. Normalized expression values of the remaining 8 out of identified 15 miRNA biomarker candidates with the highest importance scores for FSHD prediction, based on the Heier et al. dataset.

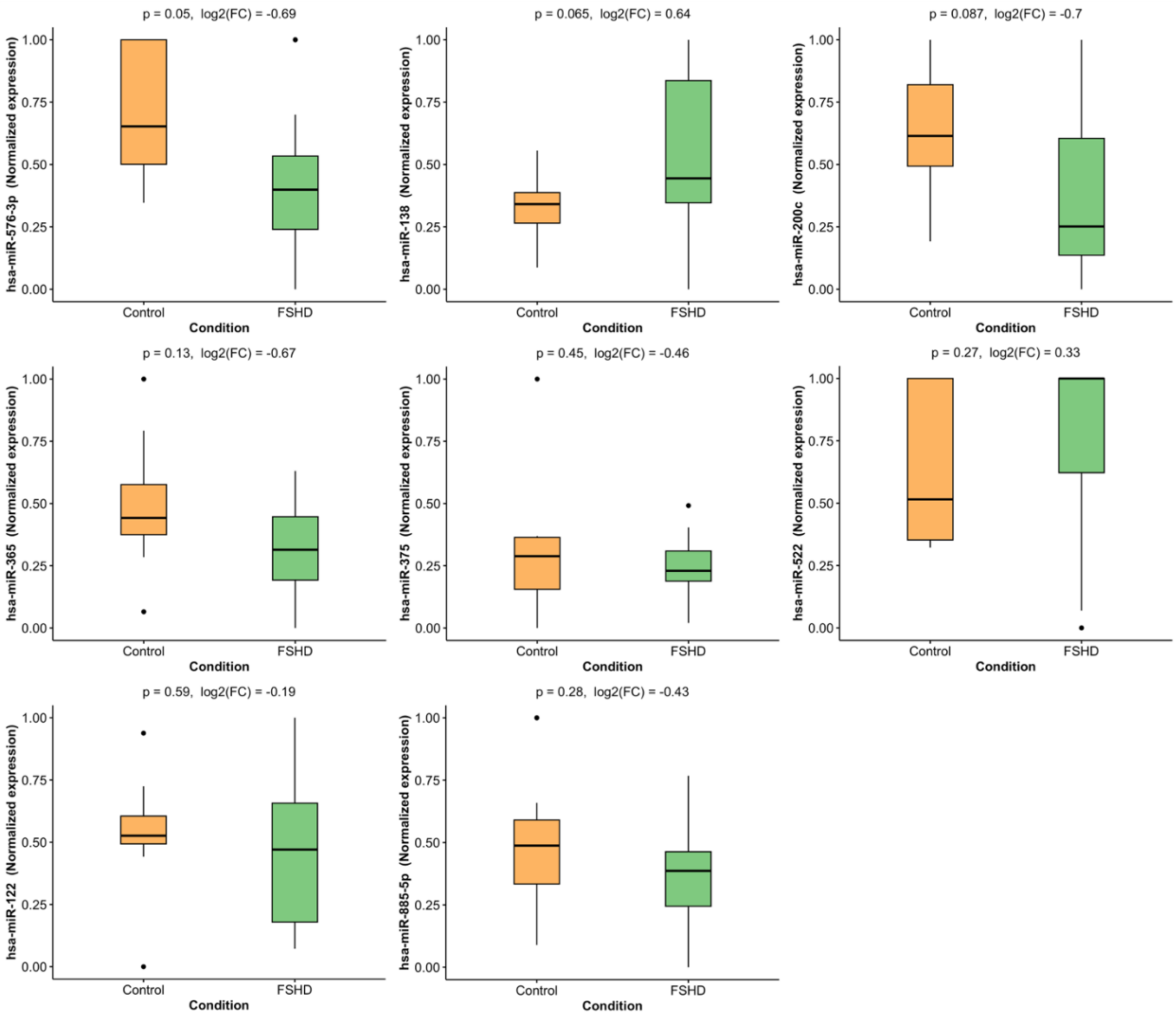

**Supplementary Material S3.** Normalized unique peptide counts of the remaining 14 out of identified 20 proteins with the highest importance for FSHD prediction, based on the Heier et al. dataset.

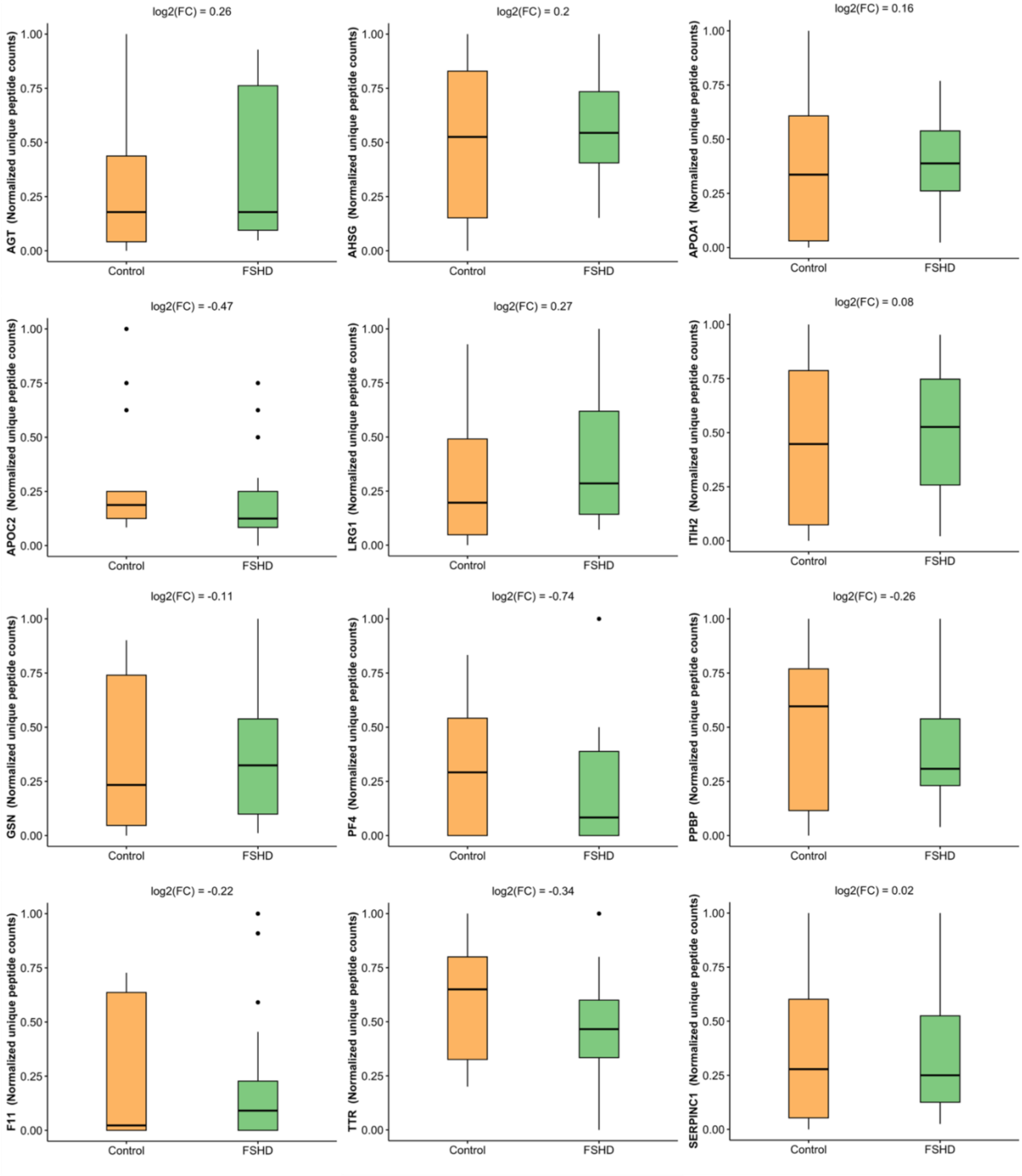

**Supplementary Material S4.** Average classification performance of the SVM and RF model with 95% confidence interval for FSHD prediction using biomarker sets, evaluated via five-fold cross-validation.

| | Metric | ML (Our) | Corasolla et al. | Statland et al. | Wong et al. | Petek et al. |
|---|---|---|---|---|---|---|
| SVM | Accuracy | 0.636±0.087 | 0.636±0.140 | 0.614±0.174 | 0.614±0.174 | 0.614±0.174 |
| | F1-score | 0.737±0.099 | 0.759±0.112 | 0.753±0.137 | 0.753±0.137 | 0.753±0.137 |
| | Precision | 0.689±0.244 | 0.642±0.185 | 0.614±0.174 | 0.614±0.174 | 0.614±0.174 |
| | Recall | 0.900±0.278 | 0.967±0.093 | 1.000±0.000 | 1.000±0.000 | 1.000±0.000 |
| RF | Accuracy | 0.761±0.273 | 0.733±0.195 | 0.686±0.221 | 0.614±0.233 | 0.639±0.156 |
| | F1-score | 0.780±0.269 | 0.780±0.170 | 0.748±0.191 | 0.718±0.174 | 0.735±0.140 |
| | Precision | 0.827±0.254 | 0.780±0.179 | 0.715±0.173 | 0.652±0.179 | 0.665±0.174 |
| | Recall | 0.817±0.361 | 0.810±0.257 | 0.810±0.257 | 0.817±0.212 | 0.850±0.185 |

**Supplementary Material S5.** FSHD prediction performance across different numbers of biomarkers using proteomics, miRNA, and multi-omics datasets. For each iteration, five-fold cross-validation was performed, progressively eliminating the least important feature for SVM, RF, and LR models, and (a) the average accuracy, (b) precision, and (c) recall were evaluated.

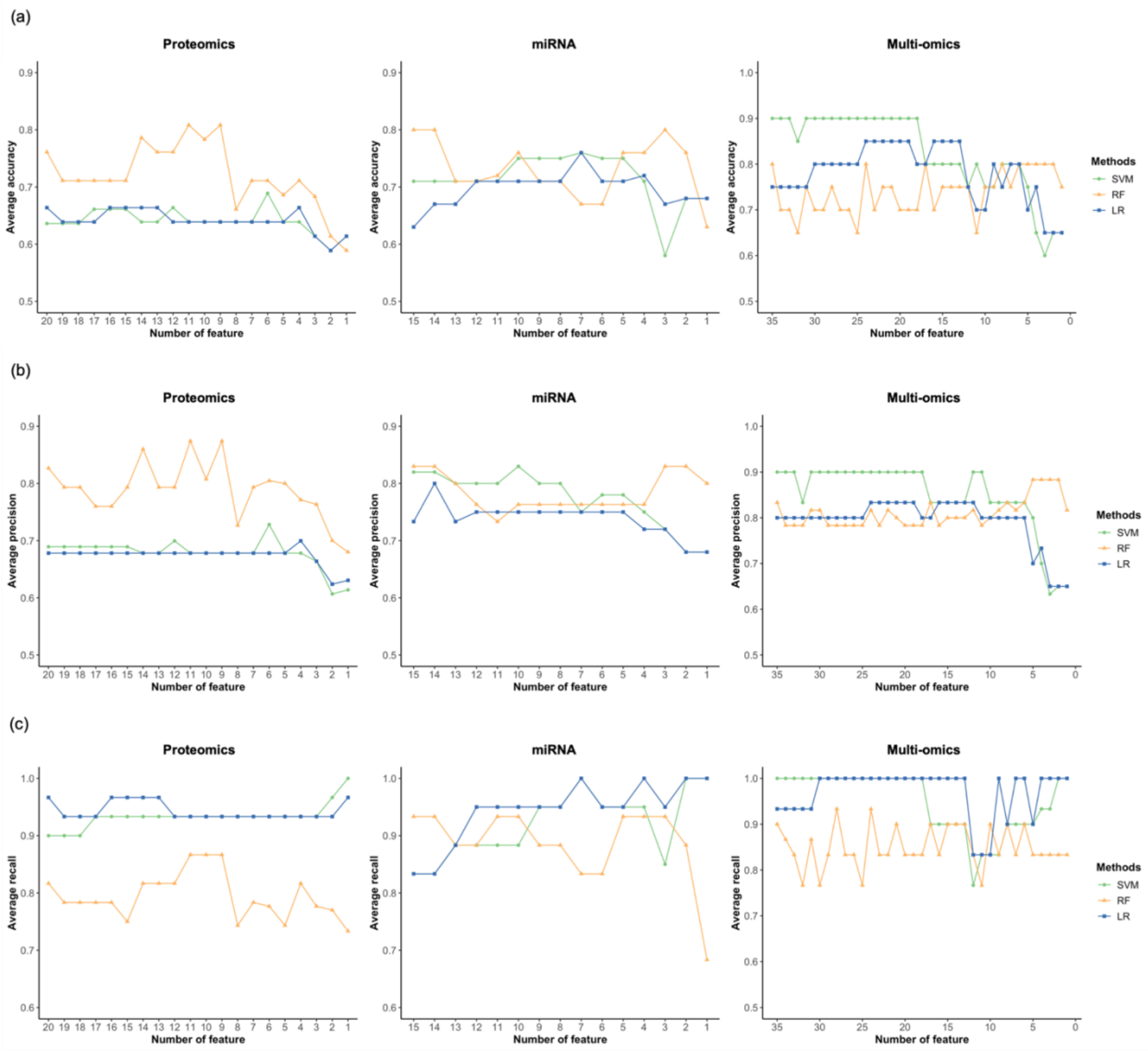
